## Supplemental Figure 1 for "Influenza A virus reassortment in mammals gives rise to genetically distinct within-host sub-populations"

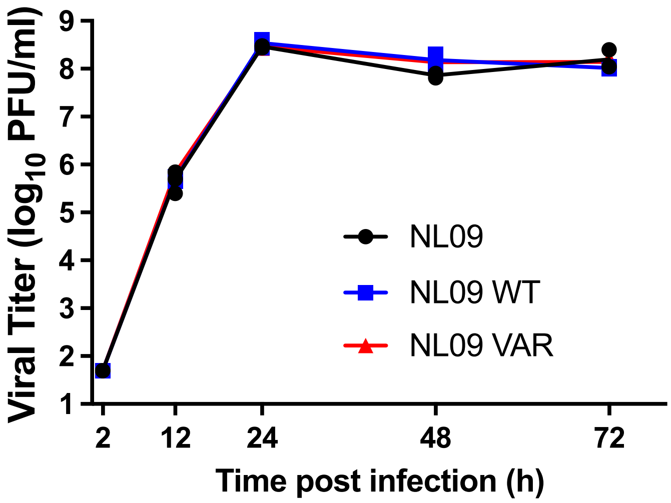


**Supplementary Fig. 1. NL09 WT and NL09 VAR show comparable replication in MDCK cells.** Viral replication from low MOI was monitored over time in three replicate culture dishes. Mean and standard deviation are plotted.
