## Supplemental Figure 2 for "Influenza A virus reassortment in mammals gives rise to genetically distinct within-host sub-populations"

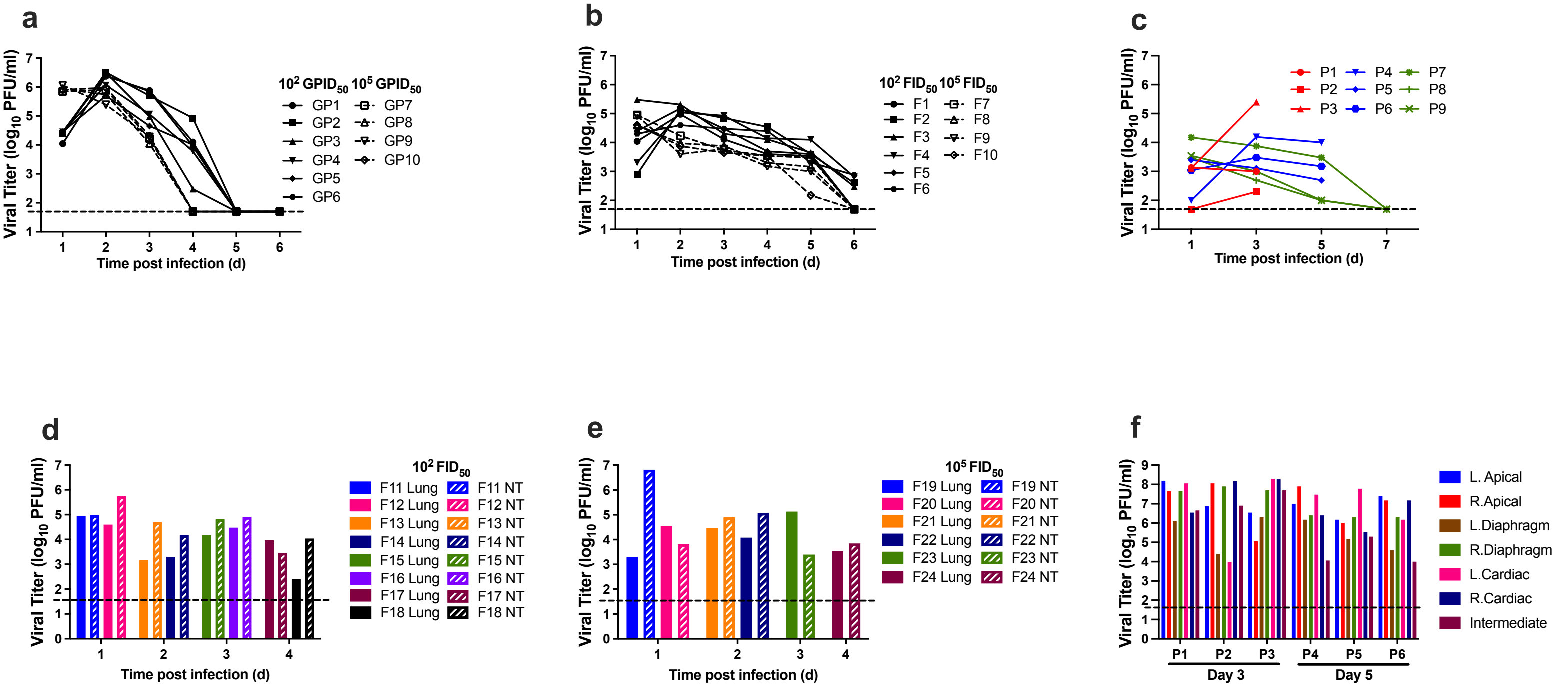


**Supplementary Fig. 2. Viral loads in infected guinea pigs, ferrets and swine.** Viral shedding determined by plaque assay from guinea pig nasal washes (A), ferret nasal washes (B), swine nasal swabs (C), ferret nasal turbinates (D), ferret lung homogenates (E), and swine lung homogenates (F) over time are shown. The horizontal dashed line represents the limit of detection of the plaque assay (50 PFU/mL).
