## Supplemental Figure 3 for "Influenza A virus reassortment in mammals gives rise to genetically distinct within-host sub-populations"

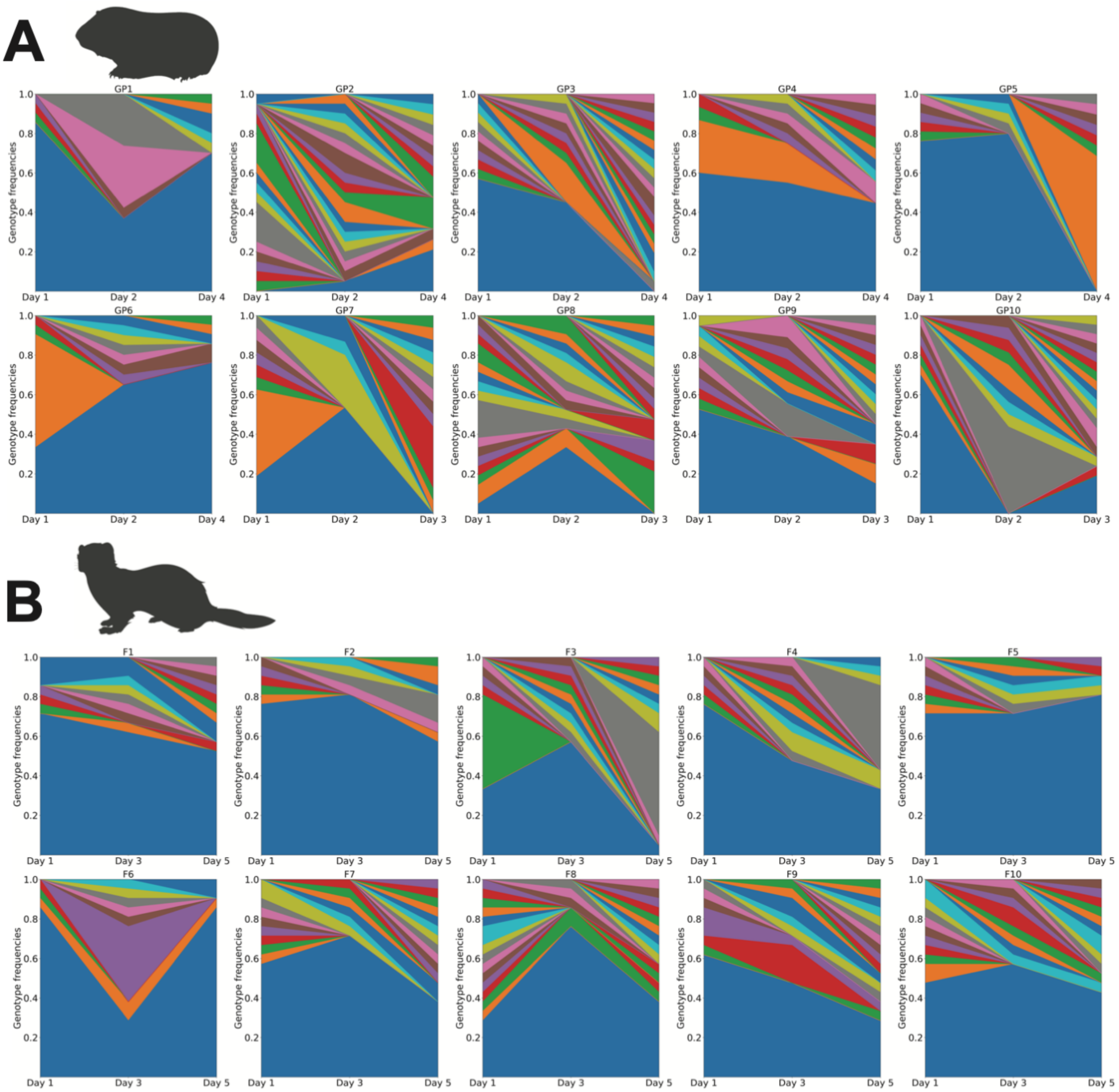


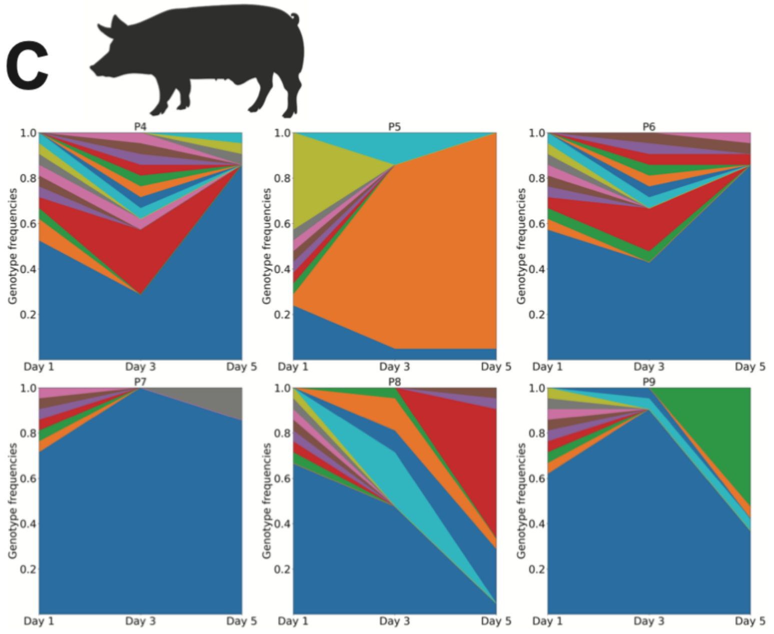


**Supplementary Fig. 3. Viral genotypic diversity generated through reassortment in the mammalian nasal tract.** (Corresponds to Figure 1) Stacked plots showing frequencies of unique genotypes detected in guinea pigs (A), ferrets (B) or swine (C) over time. The sections in blue and orange represent the WT and VAR parental genotypes, respectively
