## Supplemental Figure 4 for "Influenza A virus reassortment in mammals gives rise to genetically distinct within-host sub-populations"

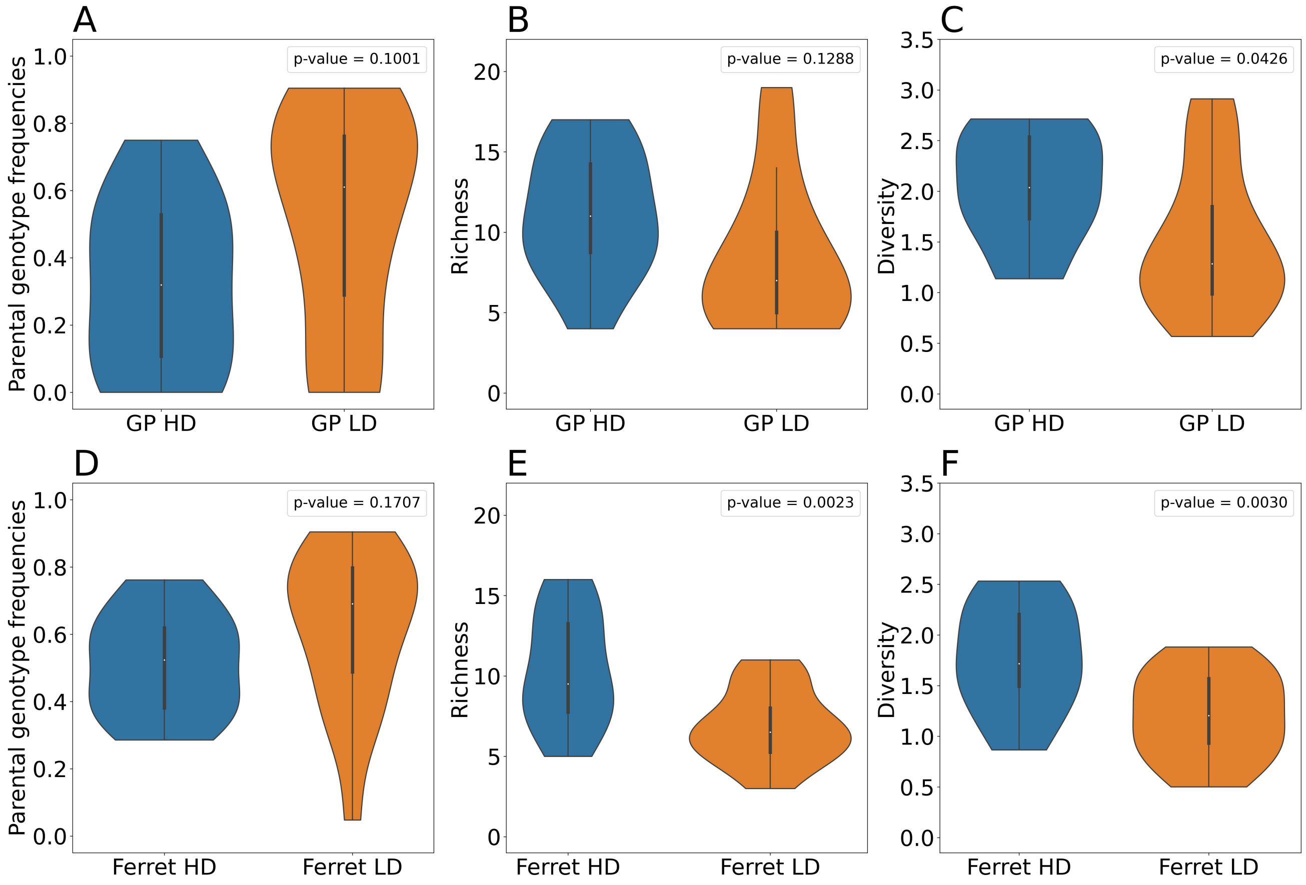


**Supplementary Fig. 4. Effects of inoculation dose on viral diversity generated through reassortment in the nasal tract.** (Corresponds to Figure 1) Results from guinea pigs are shown in panels A–C, ferrets in panels D–F. HD indicates high dose ($1x10^5$ $ID_{50}$) and LD indicates low dose ($1x10^2$ $ID_{50}$) groups. The distribution of parental genotype frequencies (A, D), richness (B, E) and diversity (C, F) across all time points in each species is shown with violin plots. P values were determined by ANOVA.
