## Supplemental Figure 5 for "Influenza A virus reassortment in mammals gives rise to genetically distinct within-host sub-populations"

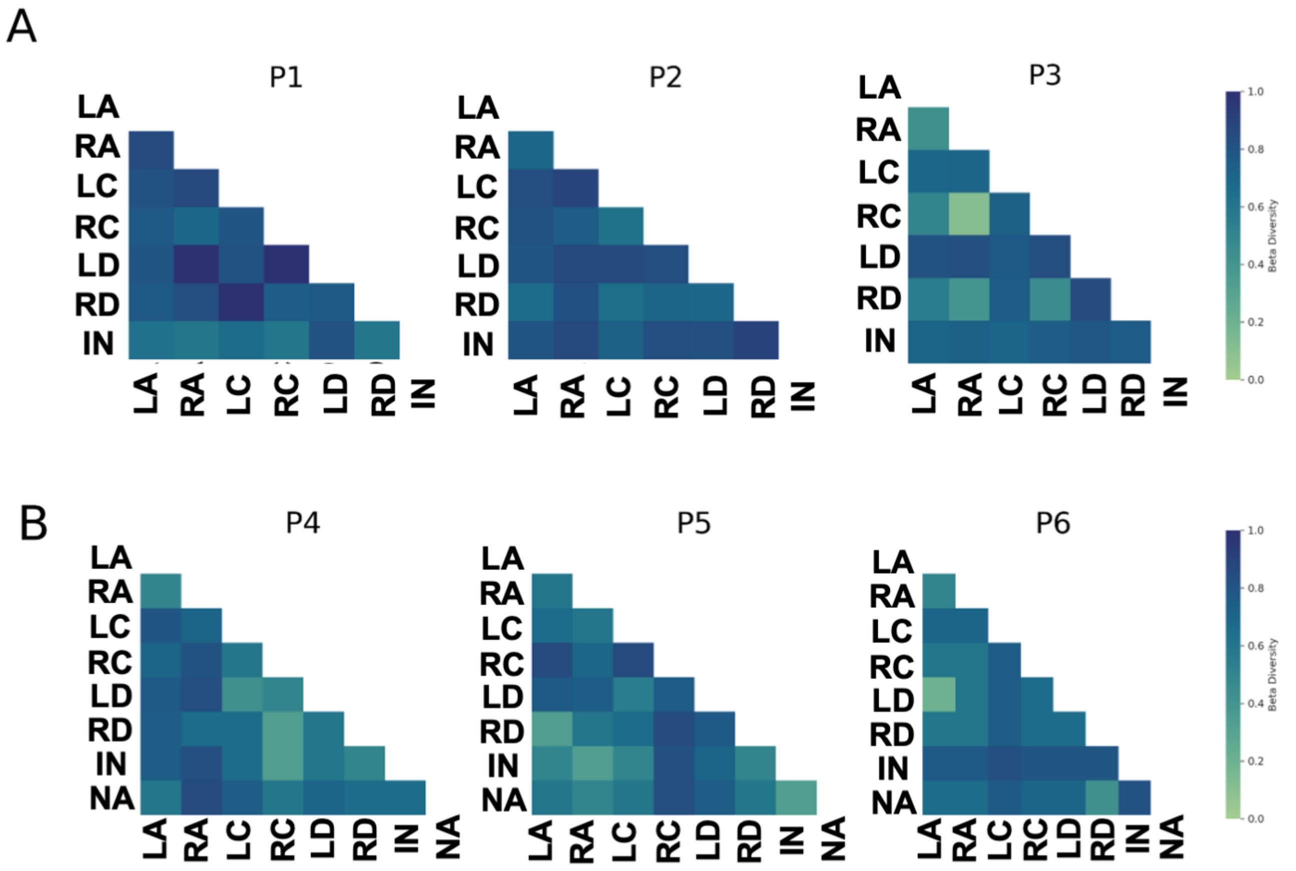


**Supplementary Fig. 5. Reassortant viral populations show extensive compartmentalization within the swine respiratory tract.** (Corresponds to Figure 5) Heat maps showing beta diversity of viral populations in the lung lobes and nasal tract of individual pigs. Pig ID number is indicated above each matrix. (A) Tissues were extracted on day 3 post-inoculation. (B) Tissues were extracted on day 5 post-inoculation. The tissue sites are abbreviated as NA-Nasal; LA- Left Apical; RA- Right Apical; LC-Left Cardiac; RC-Right Cardiac; LD-Left Diaphragmatic; RD-Right Diaphragmatic; IN-Intermediate.
