## Supplemental Figure 6A for "Influenza A virus reassortment in mammals gives rise to genetically distinct within-host sub-populations"

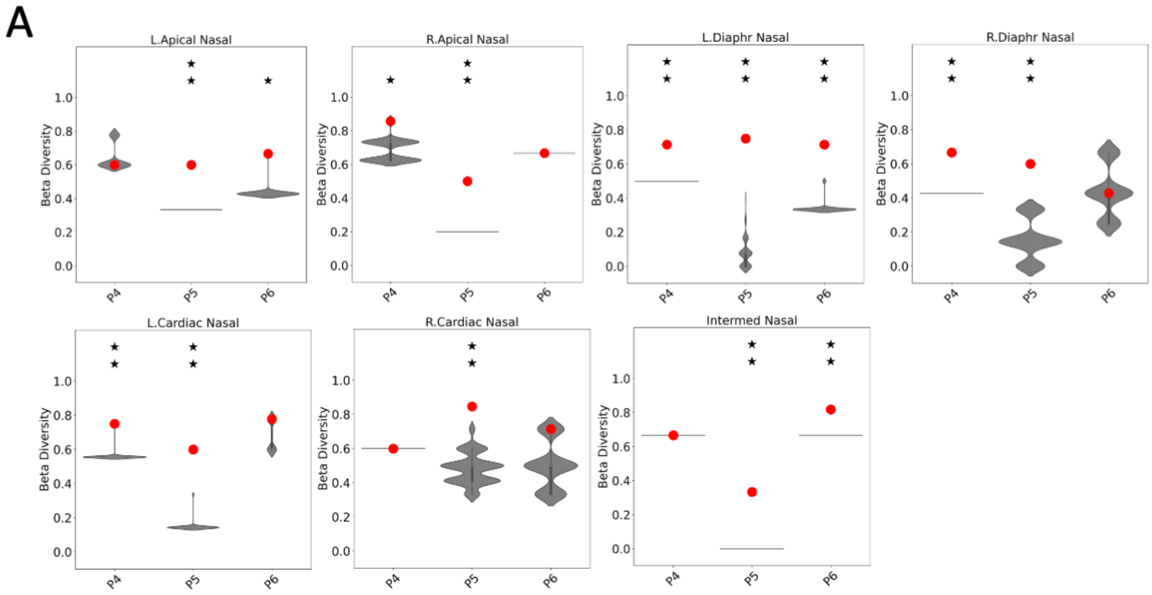


**Supplementary Fig. 6. Reassortant viral populations show extensive compartmentalization within the swine respiratory tract.** (Corresponds to Figure 5) Normalized beta diversity between each lung lobe and nasal tract (A) and between lung lobes (B) are plotted with observed results (colored points) overlaid on the distribution of simulated data (gray violins). One star indicates that observed data is above the 95$^{th}$ percentile of the distribution; two stars indicates that observed data is above the 99$^{th}$ percentile.
