## Supplementary figures and images for "Influenza A virus reassortment in mammals gives rise to genetically distinct within-host sub-populations"

### Supplemental Figure 6B

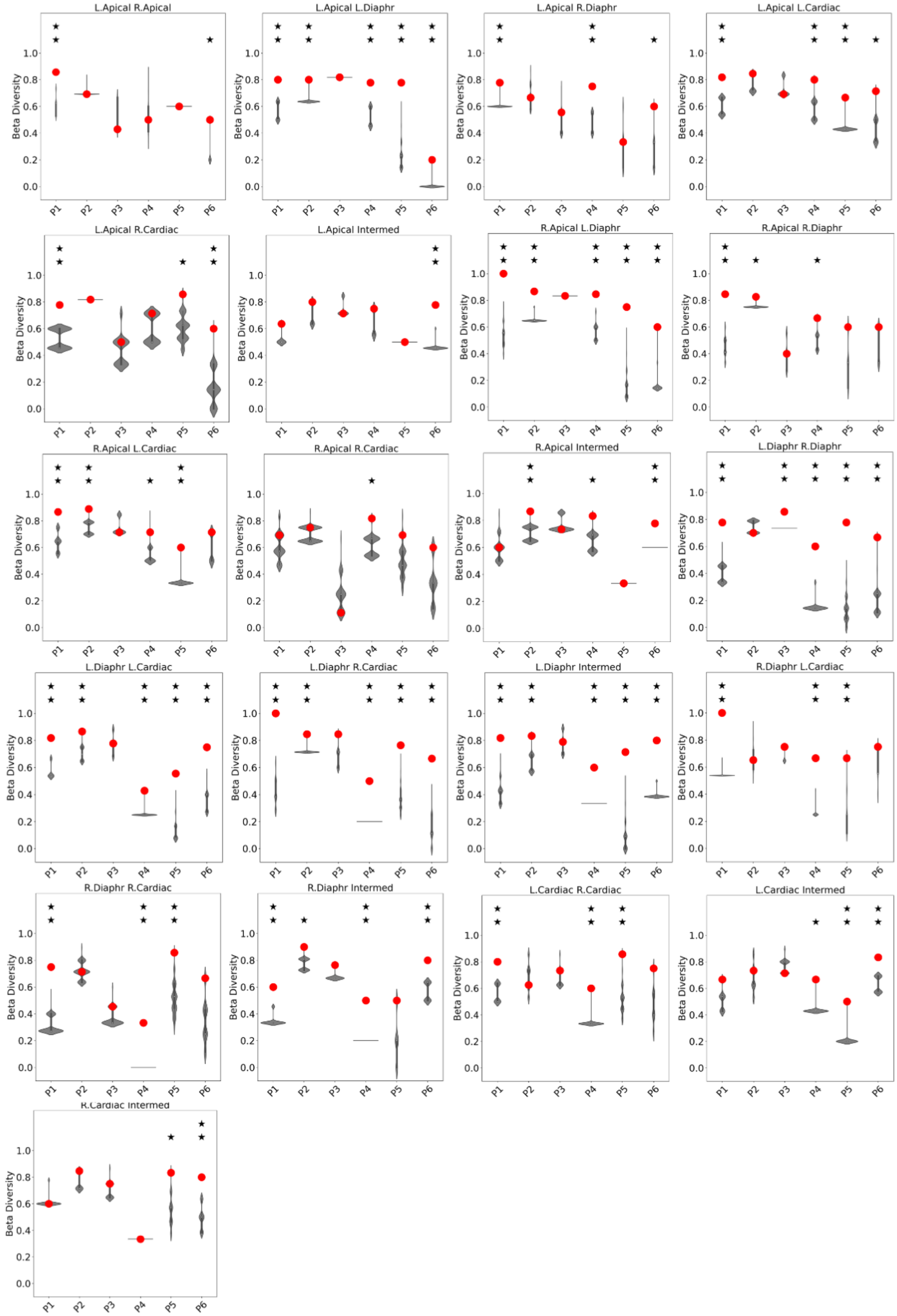

### Supplemental Figure 7-2

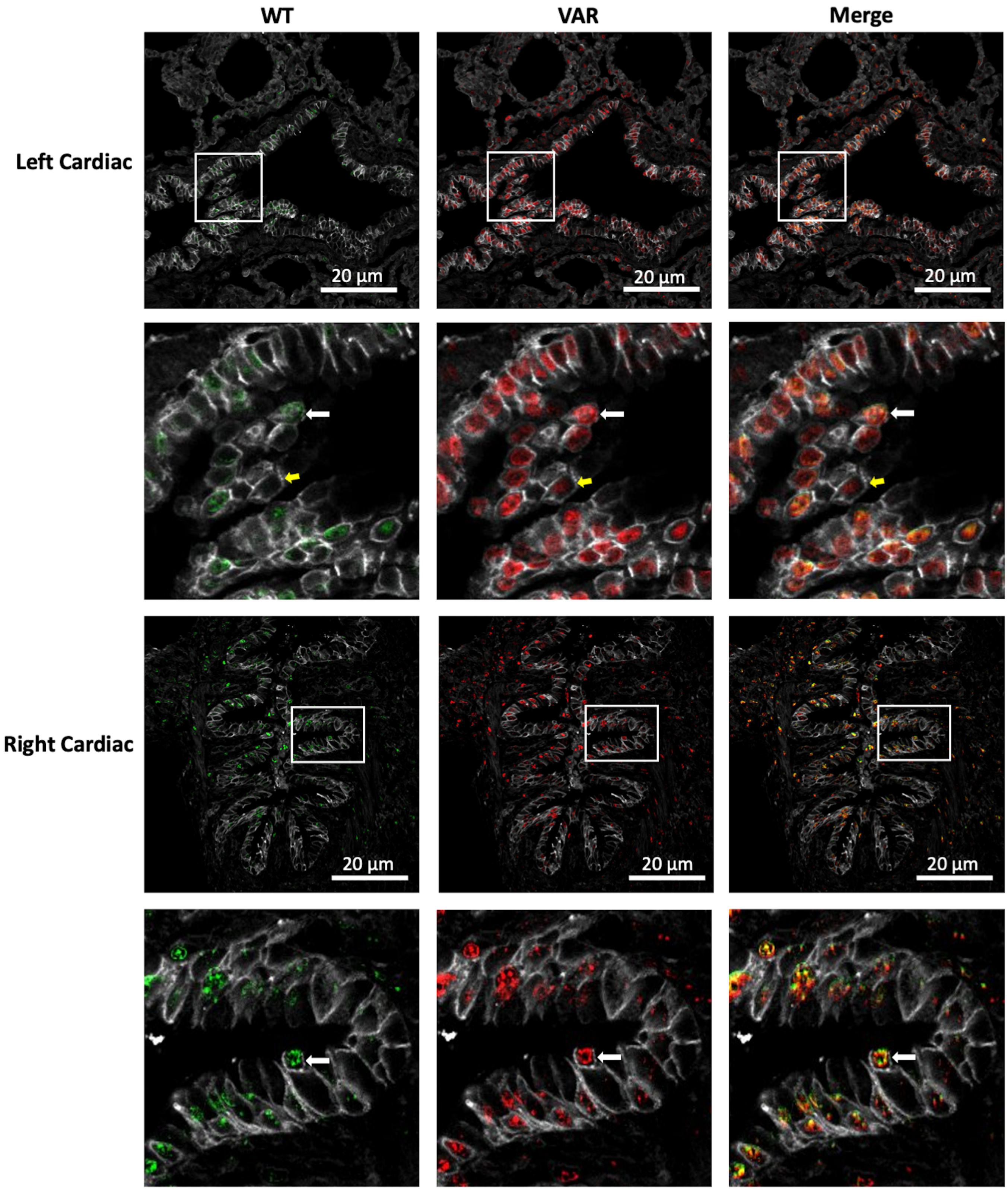

### Supplemental Figure 7-3

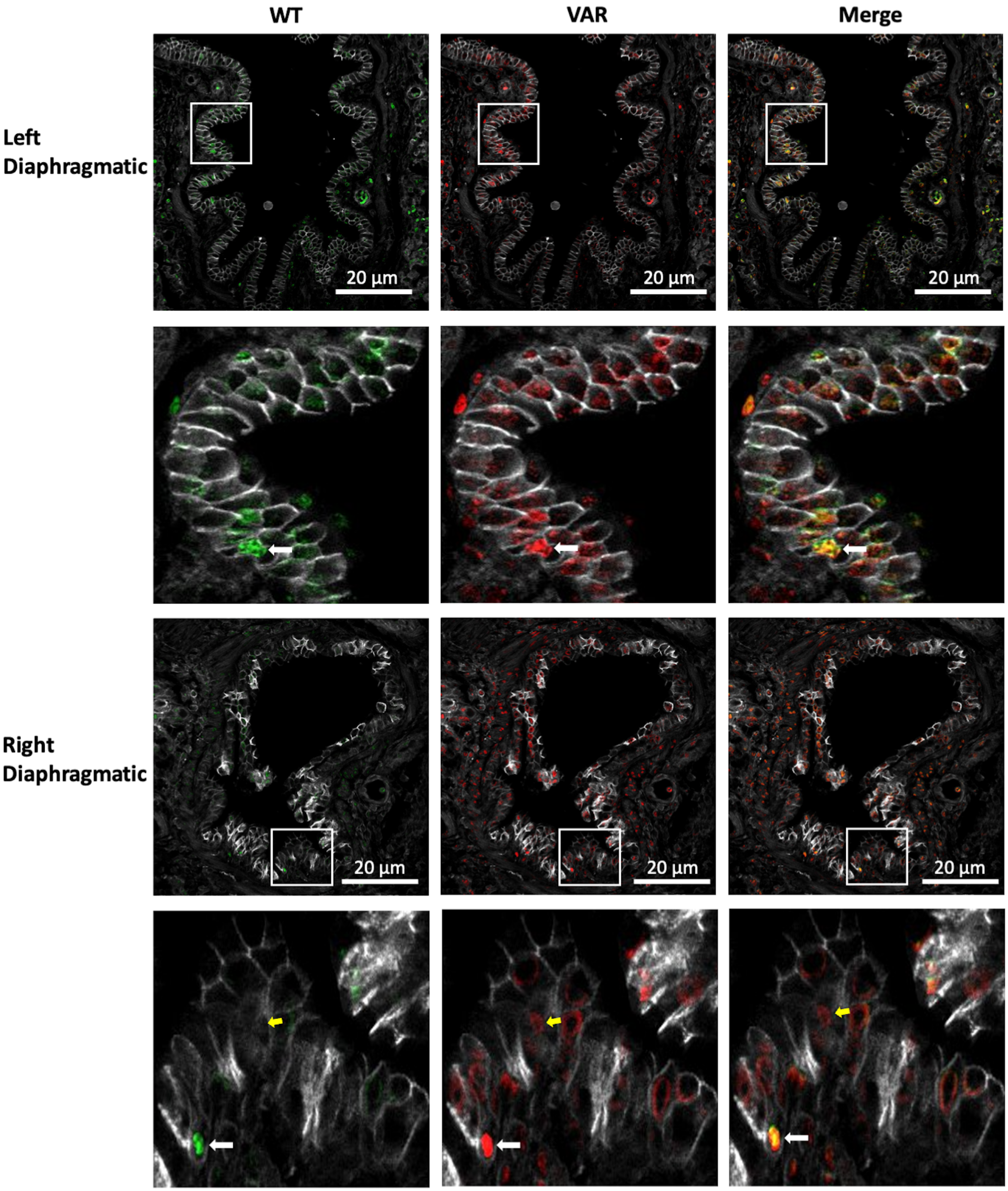

### Supplemental Figure 7-4

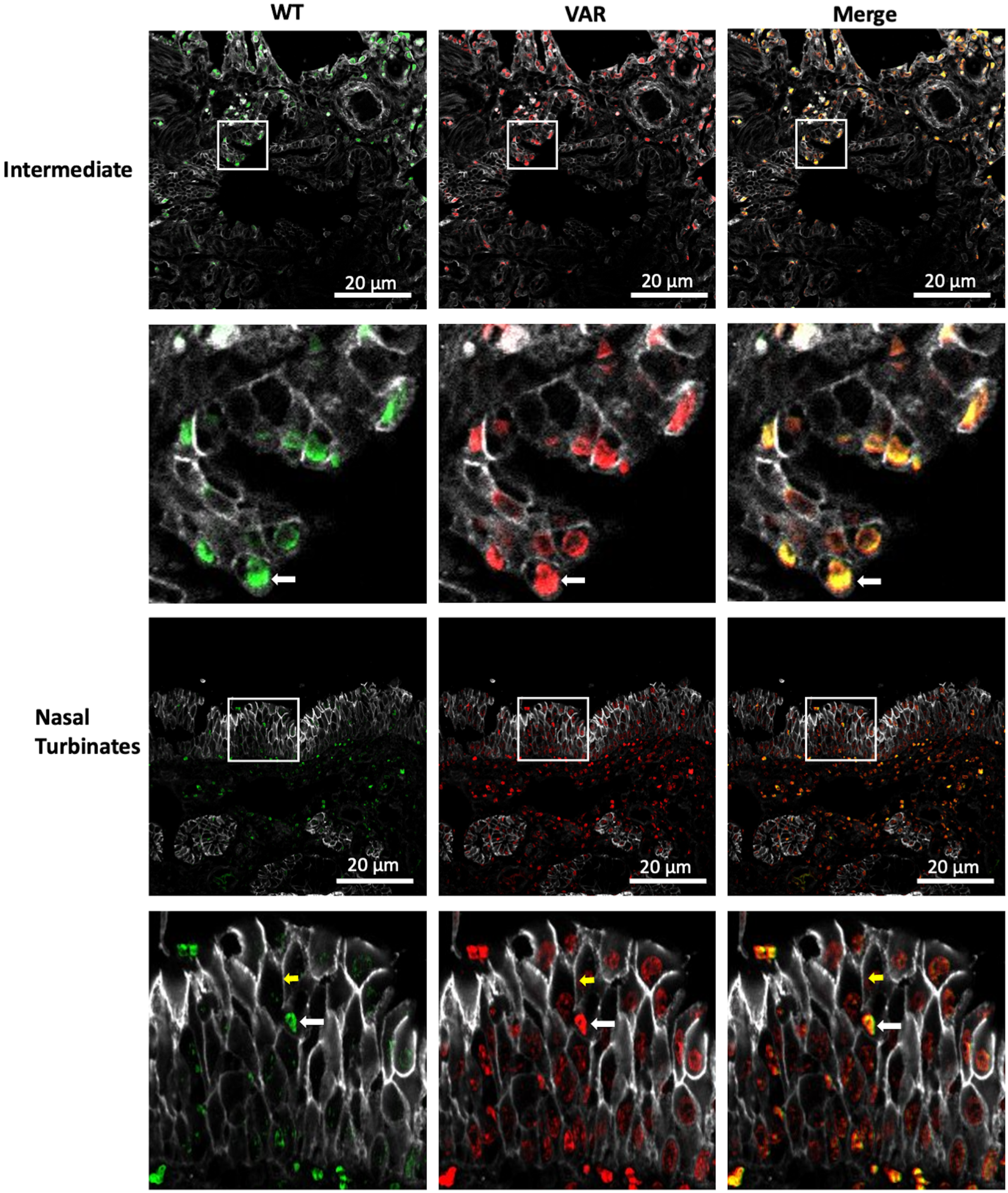

### Supplemental Figure 10-2

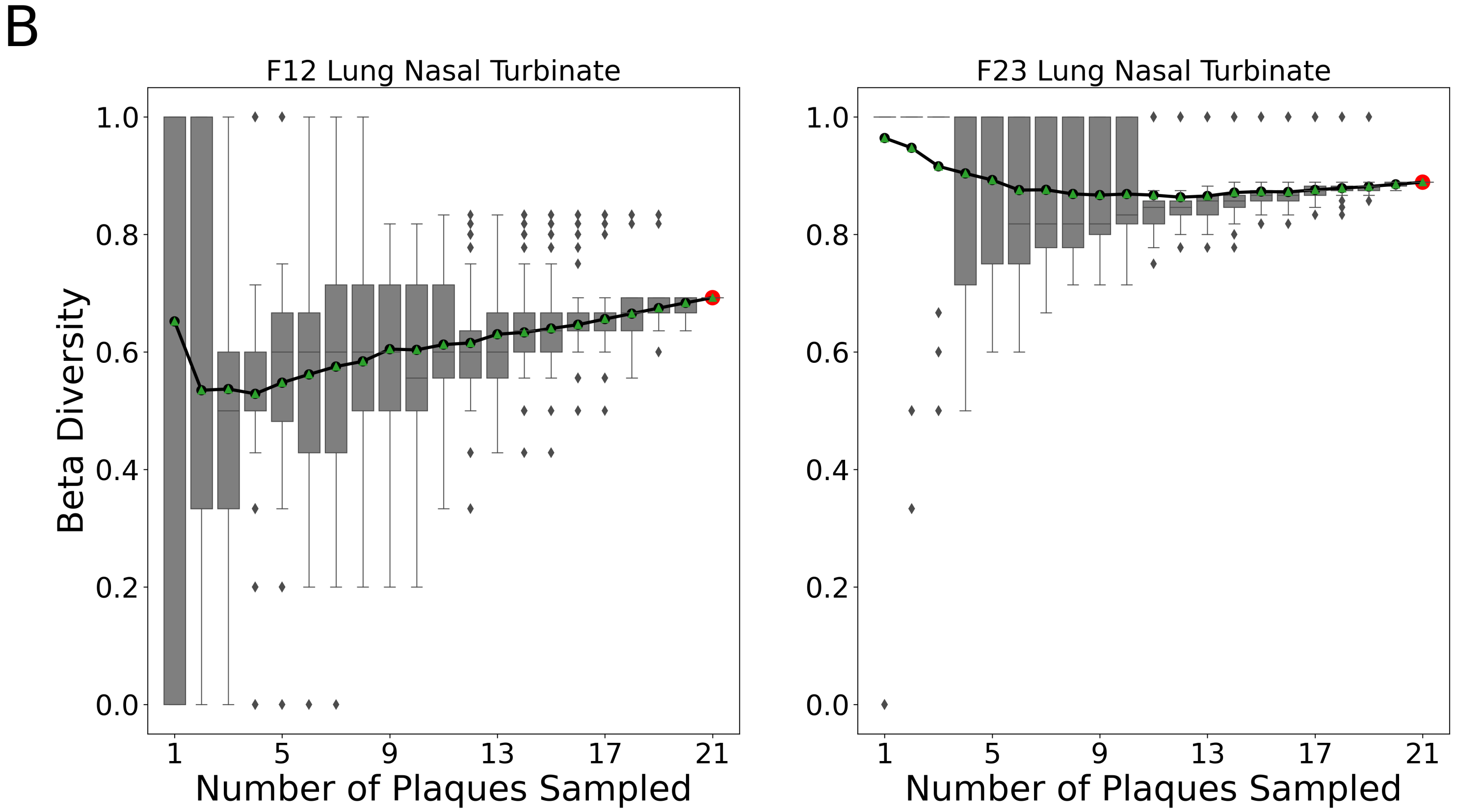
