## Supplemental Figure 7-1 for "Influenza A virus reassortment in mammals gives rise to genetically distinct within-host sub-populations"

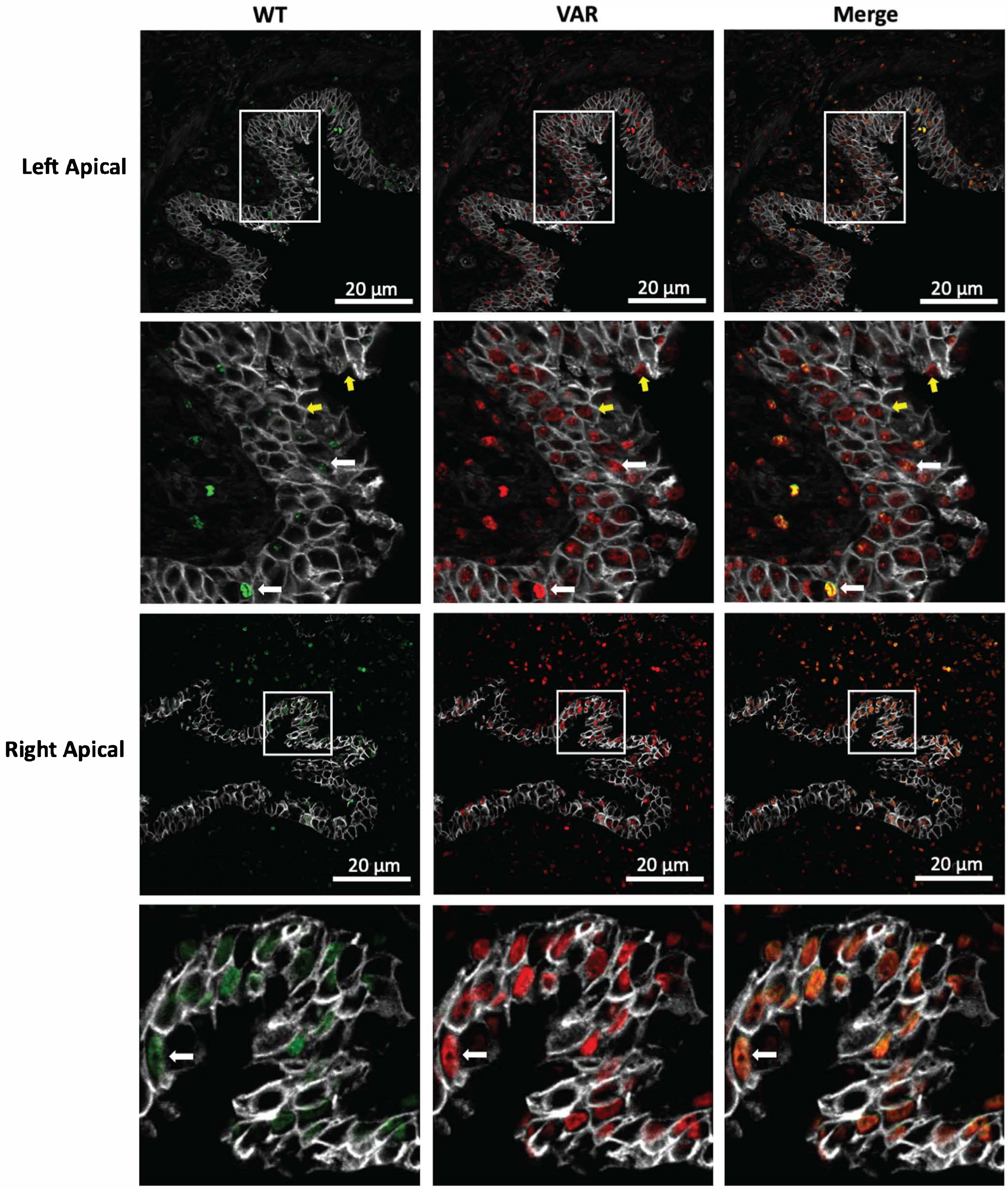


**Supplementary Fig. 7. Coinfection is common in the swine respiratory tract.** (Corresponds to Figure 5) Immunohistochemistry images of lung tissue sections stained for WT (green) and VAR (red) viruses at day 3 post-inoculation. The lung lobe sampled is indicated at the left. Gray staining marks epithelial cell borders. Yellow coloring in merged images indicates the presence of both WT and VAR HA antigens in the same cell. Zoomed insets are shown with white arrows indicating co-infected cells and yellow arrows indicating singly infected cells.
