## Supplemental Figure 8 for "Influenza A virus reassortment in mammals gives rise to genetically distinct within-host sub-populations"

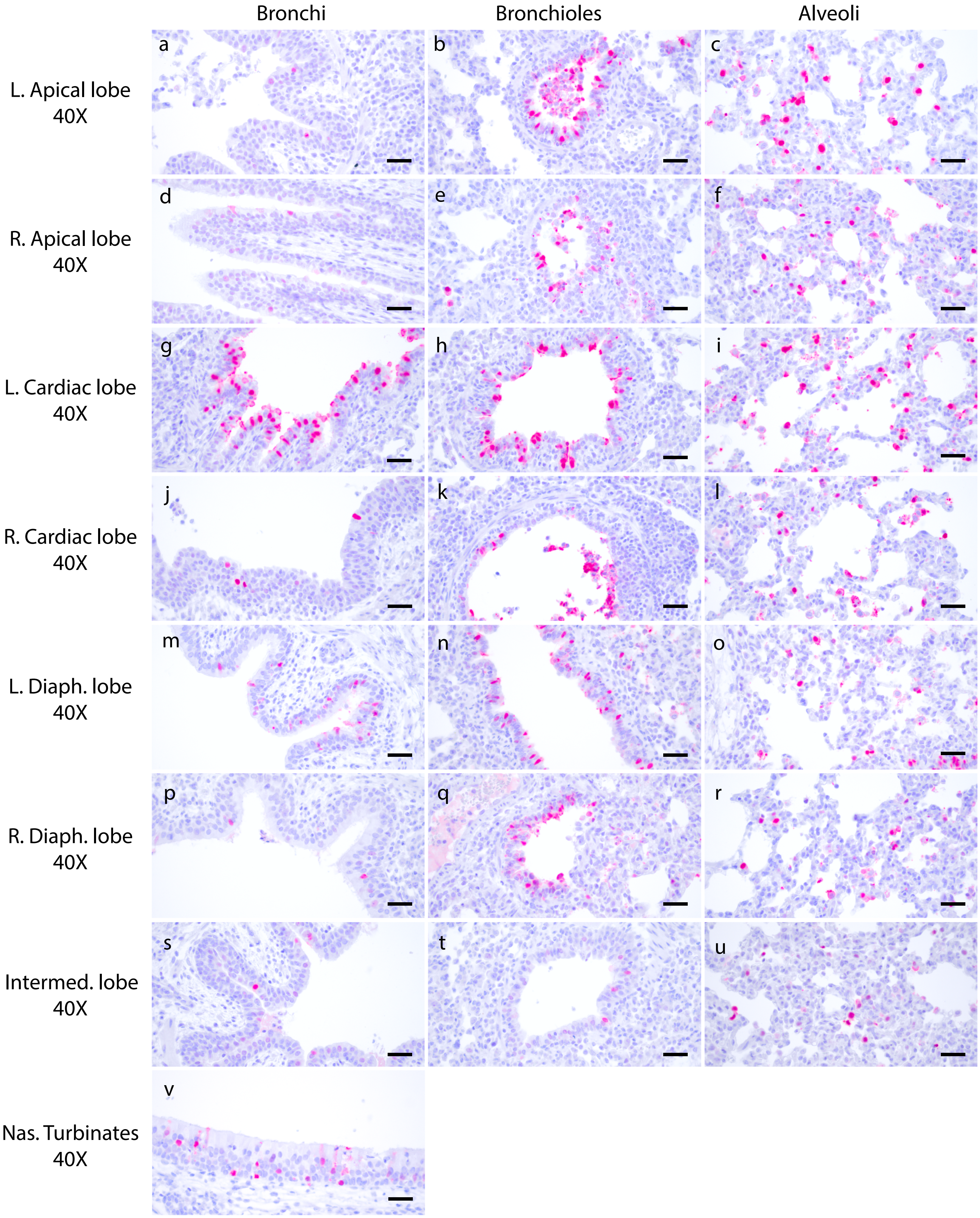


**Supplementary Fig. 8. Identification of infected cell types in swine respiratory tract.** (Corresponds to Figure 5). Tissue sections were stained for viral nucleoprotein (red) and counterstained with hematoxylin and eosin. Scale bars are 20 μm.
