## Supplemental Figure 9 for "Influenza A virus reassortment in mammals gives rise to genetically distinct within-host sub-populations"

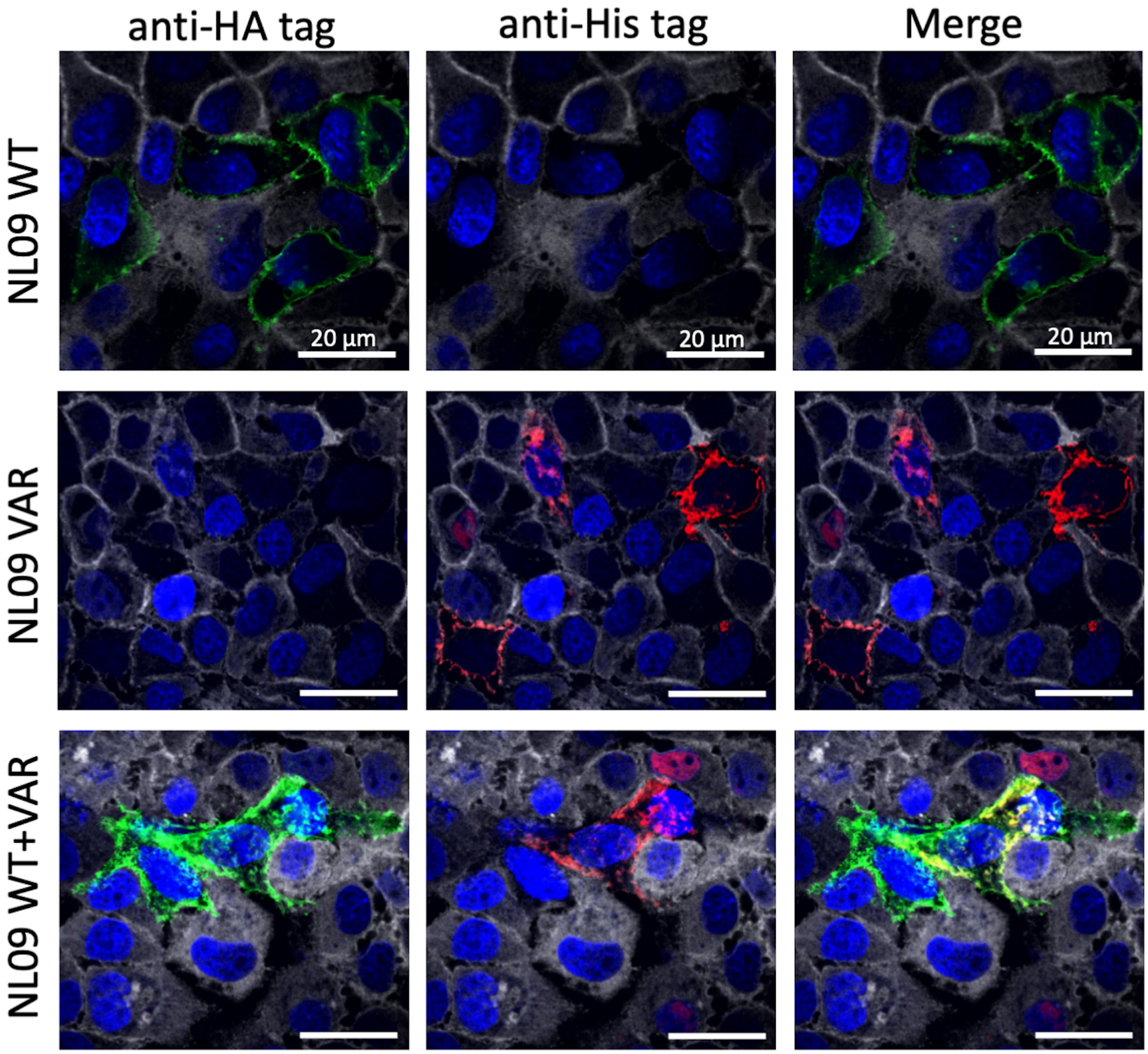


**Supplementary Fig. 9. Validation of the antibodies used for immunohistochemistry.** Immunofluorescence images of MDCK cells stained for WT (green) and VAR (red) viruses at 24 h post infection. Gray staining marks epithelial cell borders. Yellow coloring in merged images indicates the presence of both WT and VAR HA antigens in the same cell.
