## Supplemental Figure 10-1 for "Influenza A virus reassortment in mammals gives rise to genetically distinct within-host sub-populations"

# A

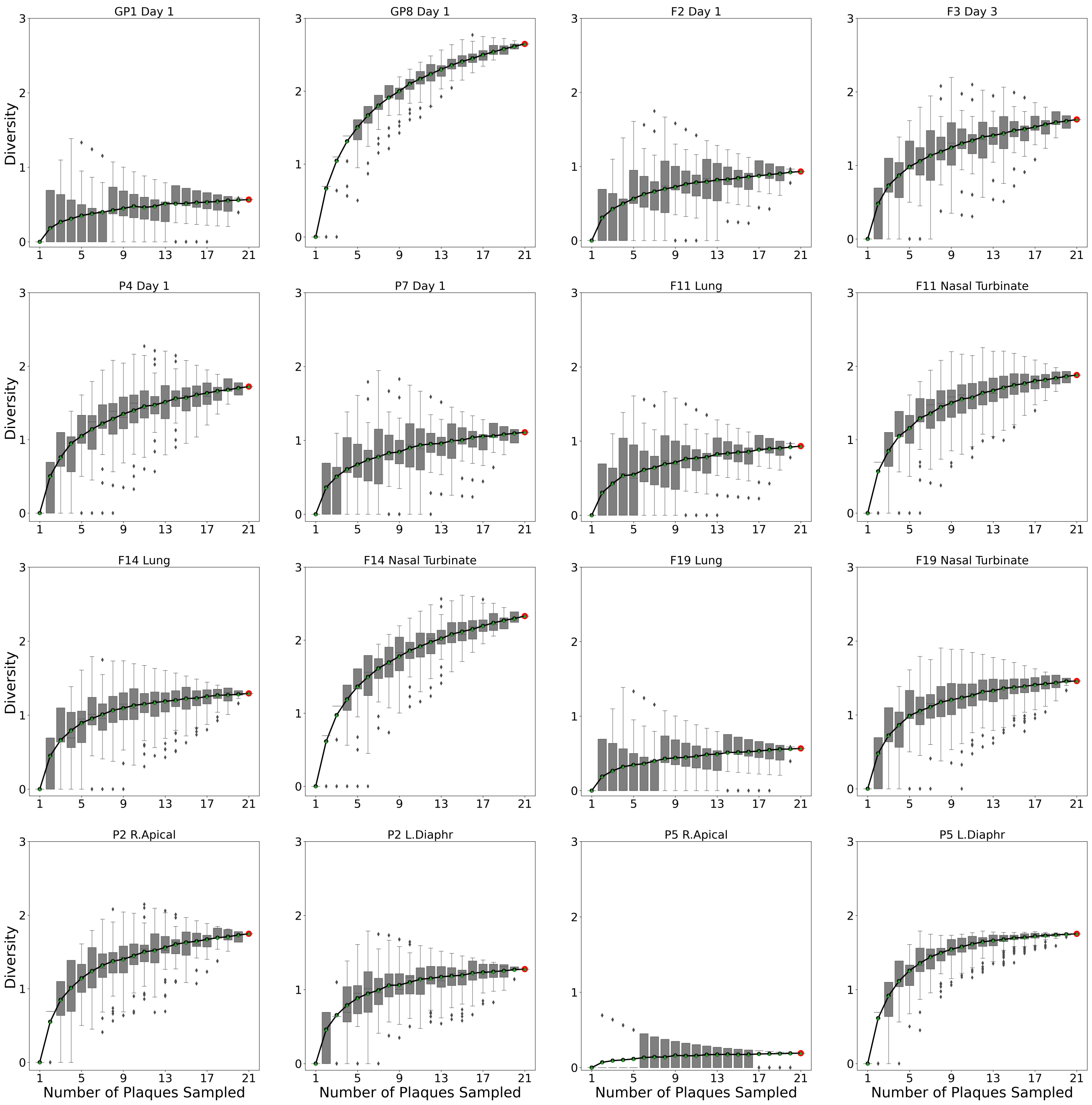


**Supplementary Fig. 10. Evaluation of the sensitivities of diversity and beta diversity to the number of viral plaques analyzed.** The impact of the number of plaques analyzed on the detection of diversity in guinea pigs, pigs and ferrets (A), beta diversity in ferrets (B) and beta diversity in pigs (C) was assessed by sub-sampling from our experimental dataset with 1000 replicate simulations performed at each number of plaques. Representative samples were analyzed and the sample is indicated above each facet. Points with the green triangles inside indicate the mean. Boxes represent the first and third quartiles, with the middle line representing the median. Whiskers show the minimum and maximum of the data with the outliers omitted outside of two standard deviations (plotted by the diamonds). The red data point shows the observed result reported in the main figures.
